# BSG.S regulates ionocyte radial intercalation and function in the developing *Xenopus* mucociliary epithelium

**DOI:** 10.64898/2026.10.30.763153

**Authors:** Camila Garcia, Hyun-Kyung Lee, Francesco Brusco, Yoo-Seok Hwang, Francisco Pereira Lobo, Ira O. Daar, Jaeho Yoon

## Abstract

Mucociliary epithelia (MCE) line multiple vertebrate organs, including the mammalian airways, and rely on coordinated interactions among specialized cell types to clear pathogens and maintain epithelial homeostasis. Proper MCE function requires the appropriate specification and spatial organization of ionocytes, multiciliated cells, secretory cells, and basal cells. Ionocytes regulate ion transport and pH homeostasis, yet the mechanisms governing their integration and function within the developing epithelium remain incompletely understood. Basigin (BSG), also known as CD147 or EMMPRIN, is a conserved transmembrane glycoprotein implicated in cell migration, adhesion, and signaling. However, its role in mucociliary epithelial development remains unclear. Here, we show that the short isoform of BSG (BSG.S) is expressed in ionocytes of the *Xenopus laevis* MCE. Our findings demonstrate that BSG.S depletion impairs ionocyte radial intercalation, alters intracellular pH, and disrupts epithelial organization. Furthermore, BSG.S loss alters epithelial cell-type composition, decreasing the number of basal cells and increasing the number of multiciliated cells. These findings suggest BSG.S as a regulator of ionocyte integration, epithelial organization, and cell-type composition during mucociliary epithelial development.

## Introduction

The mucociliary epithelium (MCE) forms a protective interface between the external environment and underlying tissues in organs such as the respiratory tract (*1, 2*). Its protective function depends on the coordinated activity of specialized epithelial cell populations. Secretory cells, including goblet cells, produce mucus that traps particles and pathogens, while multiciliated cells (MCCs) propel mucus through the beating of motile cilia. Basal cells support epithelial maintenance and renewal, whereas ionocytes regulate ion transport and acid–base homeostasis. Proper tissue function therefore depends on both the composition of these cell populations and their organization within the epithelium (*3, 4*). This functional interdependence is demonstrated by the *Xenopus* embryonic epidermis, where ionocyte depletion impairs cilia development and beating in neighboring MCCs (*5*).

The embryonic epidermis of *Xenopus* provides an accessible *in vivo* model for investigating how epithelial cell populations differentiate and assemble into a functional MCE (*6*). It shares major cell populations and aspects of their molecular regulation with mammalian mucociliary epithelia, while the embryo’s external development facilitates live imaging and experimental manipulation (*7*). The epidermis consists of an outer layer and an underlying sensorial layer containing multipotent progenitor cells. During neurulation, specified ionocyte and MCC precursors begin to undergo radial intercalation, moving toward the surface and inserting between existing superficial cells (*6*). This process coordinates changes in cell position and morphology with the establishment of cell–cell contacts and cell-type-specific differentiation, resulting in the formation of an organized mucociliary epithelium.

Ionocyte differentiation is regulated by the transcription factor Foxi1, which promotes the expression of genes encoding specialized ion transport machinery (*8*). Ionocytes express vacuolar H - ATPases (V-ATPases), chloride/bicarbonate exchangers, and carbonic anhydrases that collectively support ion transport and pH regulation. Carbonic anhydrases catalyze the reversible interconversion of CO and H O ↔ H and HCO, providing substrates for proton pumps and bicarbonate transporters. Two major ionocyte subtypes have been identified in the *Xenopus* epidermis. Alpha ionocytes express anion exchanger 1 (AE1), encoded by slc4a1, and display apically localized V-ATPase, whereas beta ionocytes express pendrin, encoded by slc26a4, and exhibit predominantly basolateral or cytoplasmic V-ATPase localization (*9*). These distinct transporter arrangements are consistent with complementary roles in proton and bicarbonate secretion, respectively. Although regulators of ionocyte differentiation and transport machinery have been identified, the molecular mechanisms coordinating radial intercalation remain poorly understood.

Basigin (BSG), also known as CD147 or EMMPRIN, is a conserved, glycosylated, single-pass transmembrane protein of the immunoglobulin superfamily. Its common isoform contains two extracellular immunoglobulin-like domains and a short cytoplasmic tail. Through interactions with membrane-associated and extracellular partners, BSG is implicated in cell adhesion, migration, and Wnt signaling (*10–14*). These properties suggest that BSG may help coordinate changes in ionocyte physiology with the cell interactions needed for ionocytes to intercalate into the epithelium. Consistent with our hypothesized role in ionocyte development, single-cell transcriptomic profiling has identified *BSG.S* among ionocyte-associated genes in the *Xenopus* MCE. However, its function in ionocyte intercalation and physiology, and its relationship to signaling pathways that regulate epithelial cell composition, remains unknown.

Here, we investigate BSG function in the developing *Xenopus* mucociliary epidermis. We show that BSG is enriched in ionocytes and that its depletion impairs their radial intercalation. These defects are accompanied by elevated intracellular pH in both ionocytes and MCCs, as well as alterations in epithelial cell-type composition and spatial organization. Specifically, BSG depletion reduces the basal cell population while increasing the multiciliated cell population. Our findings further implicate Wnt/β- catenin signaling in the epithelial defects associated with BSG loss. These findings identify BSG as a regulator of ionocyte intercalation, cell-type composition, and spatial organization in the mucociliary epithelium.

## Results

### BSG.S is enriched in ionocytes

To investigate the molecular features of ionocyte development, we re-analyzed a published single-cell RNA-sequencing dataset of the developing *Xenopus* MCE (*15*). Ionocytes were identified by the expression of established markers, including the transcription factor Foxi1 and the V-ATPase subunit atp6v1b1. This analysis identified *BSG.S* expression as specifically enriched in the ionocyte population relative to other epithelial cell populations (Fig. 1a). To validate this expression pattern, we performed hybridization chain reaction (HCR) *in situ hybridization*, which confirmed colocalization of *BSG.S* transcripts and Foxi1-positive ionocytes, with little to no expression in other MCE cell types (Fig. 1b).

**Figure 1.**
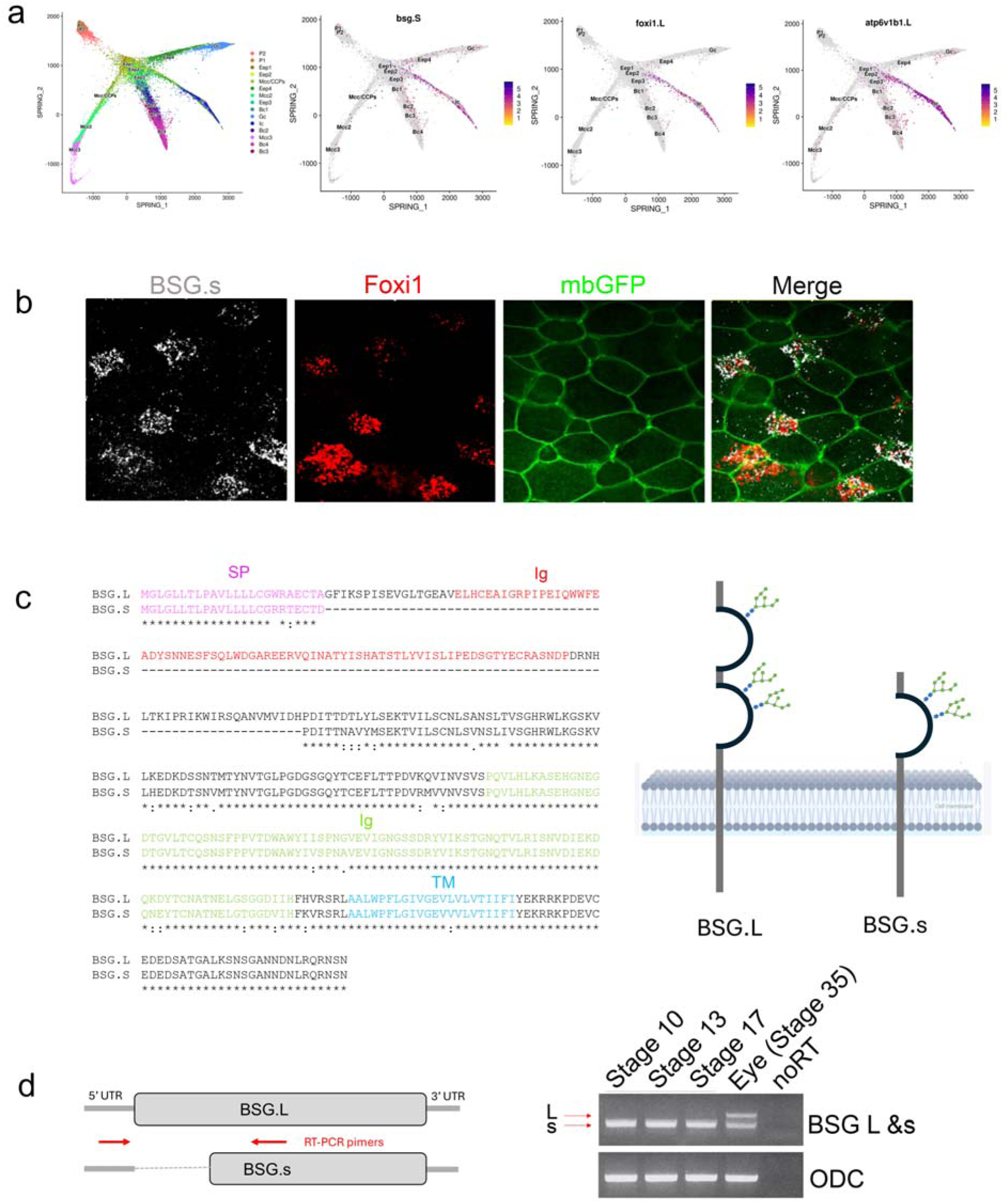
BSG.S is expressed in ionocytes of the *Xenopus laevis* mucociliary epithelium. a. Single-cell RNA sequencing data of the developing *Xenopus laevis* MCE organized by transcriptional profile and developmental stage. Feature plots depict the expression patterns of *BSG.S* as well as two ionocyte markers, *foxi1.L* and *atp6v1b1.L*. b. RNA in situ hybridization (HCR) assay showing *Foxi1* (red) and *BSG.S* (white) transcripts. Membrane- GFP (100pg) labels cell boundaries. c. Sequence alignment of two isoform of *Xenopus* BSG L and s. Structural representation of BSG.L and BSG.S protein isoforms in the *Xenopus laevis*. SP: signal peptides; Ig: immunoglobulin domain; TM: transmembrane domain. d. Schematic of RT-PCR primer-binding sites for BSG.L&S transcripts. RT-PCR analysis of BSG.L&S expression in ectodermal explants at stages 10, 13, and 17 and in eye tissue at stage 35.

BSG isoforms differ in their extracellular immunoglobulin (Ig)-like domain architecture. Four isoforms have been described in humans, while the two *Xenopus* laevis isoforms examined here comprise long and short forms with distinct extracellular domains (Fig. 1c). To determine which isoform is expressed in the developing MCE, we performed RT-PCR using primers targeting the region encoding the extracellular domain. Both long- and short-form transcripts were detected in eye tissue as demonstrated in a previous reported (*16*), whereas only the short-form transcript was detected in the developing MCE (Fig. 1d).

Our findings show that *BSG.S* expression is enriched in ionocytes during MCE development, suggesting a potential role for BSG.S in ionocyte development.

### Loss of BSG.S impairs ionocyte radial intercalation

To investigate the role of BSG.S in ionocyte development, we performed loss-of-function analysis using a BSG.S-specific morpholino oligonucleotide (MO). The MO efficiently suppressed BSG.S expression (Fig. 2a). Interestingly, BSG.S-depleted embryos exhibited epidermal abnormalities characterized by cell aggregation, darkening, and protrusions from the epidermal surface (Fig. 2b). Given the enrichment of BSG.S expression in ionocytes, we first examined whether BSG depletion altered the number of ionocytes. *In situ hybridization* analysis with antisense *Foxi1* probes showed that the number of *Foxi1*-positive cells did not differ significantly between control and BSG.S-depleted embryos. We also assessed *Foxi1* expression in an *ex vivo* system, animal-cap-derived mucociliary organoids using reverse transcription–quantitative PCR (RT-qPCR). Embryos were injected with BSG.S MO at the animal pole at the one-cell stage, and animal caps were isolated at stage 8 and cultured to the equivalent of stage 28.

**Figure 2.**
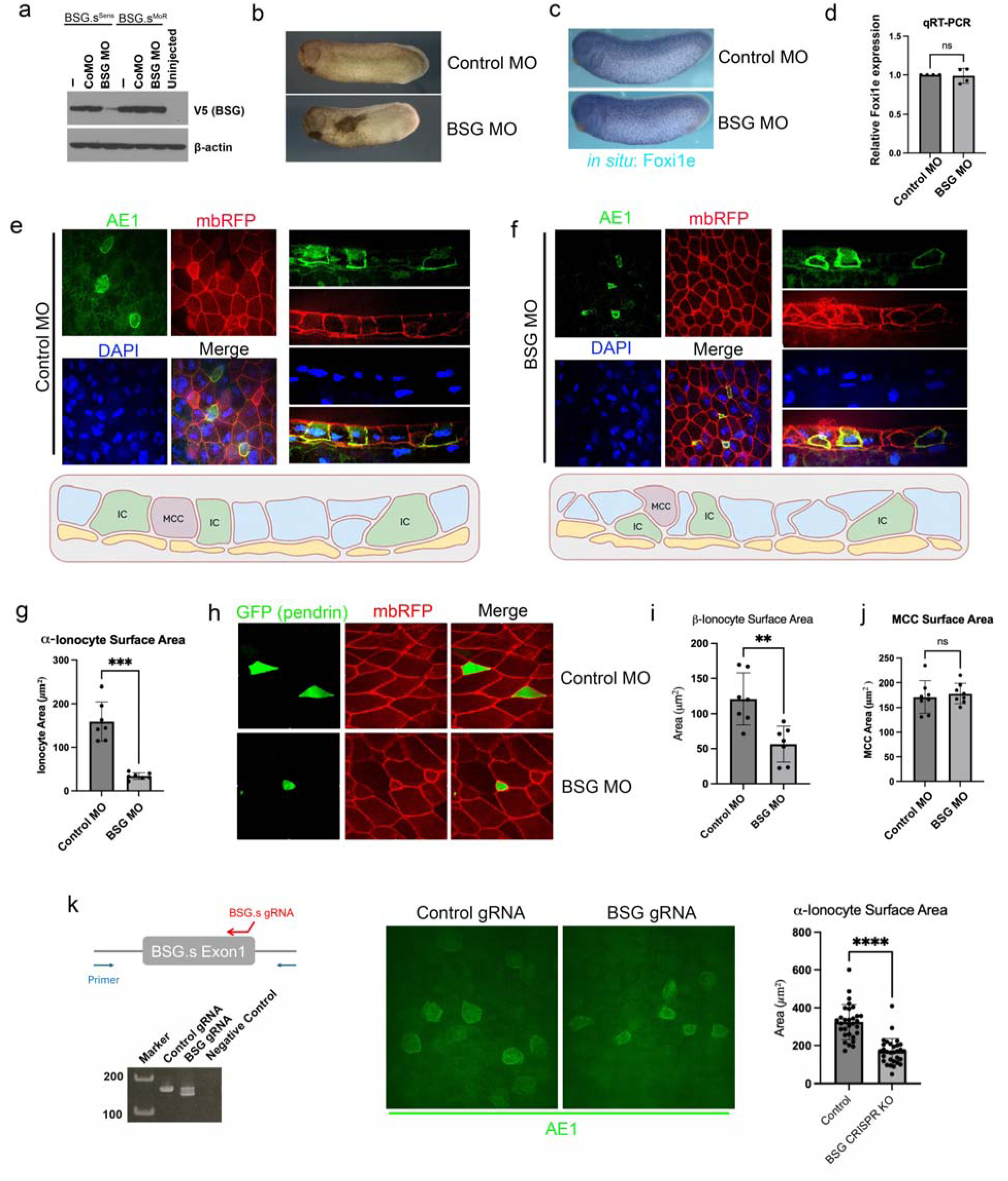
BSG.S is required for ionocyte intercalation. a. BSG.S morpholino oligos (MO). BSG.S MO targets the translational start region. Western blot analysis shows that BSG.S MO efficiently blocks exogenous BSG.S-wild type (BSG.S^Sens^) expression not morpholino resistant BSG.S (BSG.S^MoR^). b. Representative images of stage 28 *Xenopus laevis* embryos following injection of control MO or BSG.S MO (8 ng/nl) into V.1 blastomere at the 8-cell stage. c. Whole-mount *in situ* hybridization for Foxi1 (ionocyte) in control and BSG.S morphant embryos at stage 28. d. RT-qPCR analysis of Foxi1 expression in ectodermal explants (stage 28) from control and BSG.S morphant embryos revealed no statistically significant difference between groups. e & f. Immunostaining for a-ionocytes in control and BSG.S morphants (stage 28). AE1 (Green) is used for a-ionocytes marker. whole-mount *Xenopus* embryos (left) and transverse sections (right) from Control MO or BSG.S MO injected embryos. DAPI (blue) marks the nucleus; mbRFP (red) marks the cell boundaries. Schematics illustrate the organization of ionocytes in each group. g. Quantification of the apical surface area of AE1-positive a-ionocytes in control and BSG.S morphant embryos. h. Confocal images of Control and BSG.S morphant embryos (stage 28). Both groups were co-injected with a DNA reporter construct in which GFP expression is driven by the Pendrin promoter. i. Quantification of the apical surface area of Multiciliated cells in control and BSG.S morphant embryos. j. Quantification of the apical surface area of b-ionocyte (GFP positive by the Pendrin promoter) in control and BSG.S morphant embryos. k. AE1 (Green) immunostaining in BSG.S crispants. BSG.S gRNA injected embryos (F0) shows nucleotides deletion in BSG.S first exon region (left). Quantification of the apical surface area of AE1- positive a-ionocytes (right). Data are presented as mean ± [SD/SEM]. Statistical significance was determined using [paired t test]. *n* = [8] for each group. ns, not significant; \*\**p* < 0.001, \*\*\**p* < 0.0001, ****P<0.0001.

These explants differentiate into mucociliary epidermis (MCE), providing an organoid model of MCE development. Consistent with the *in situ* hybridization results, BSG.S depletion did not significantly alter *Foxi1* expression (Fig. 2d). Together, these findings are consistent with no detectable change in the abundance of ionocytes following BSG.S depletion.

Previous cancer studies have implicated BSG in cell migration and invasion through its effects on extracellular matrix (ECM) degradation (*12*). Since intercalation of ionocytes into the mature MCE may be affacted by alterations in the ECM, we examined whether BSG contributes to ionocyte morphology and radial intercalation. A previous study demonstrated that radial intercalation of multiciliated cells and ionocytes begins at stage 17 (the late neural fold stage) and is complete by stage 28 (tailbud stage) (*17*). To visualize α-ionocytes, we performed immunostaining for AE1, a chloride/bicarbonate exchanger involved in acid–base regulation. AE1-positive ionocytes in BSG.S morphants exhibited significantly smaller apical surface areas than those in control MO-injected embryos at stage 28, suggesting that the ionocytes may not have fully reached the MCE surface. Cross-sectional imaging of the MCE clearly revealed defects in ionocyte radial intercalation in BSG.S morphants (Fig. 2e-g). Similarly, GFP expressing β-ionocytes by pendrin promoter in BSG.S morphants exhibited narrower apical surface suggesting a potential intercalation defect in both α and β-ionocytes (Fig 2h-i). In contrast, the apical surface areas of multiciliated cells were unchanged, indicating the defect is specific to ionocytes (Fig. 2j). As an additional control, we also performed CRISPR-Cas12 mediated knockout of BSG.S using a guide RNA (gRNA) targeting the first exon. Genomic deletions at the target site were assessed by PCR (Fig. 2k). BSG.S knockout crispants also reduced the apical surface area of α-ionocytes, confirming the morpholino phenotype we observed in Fig. 2f.

Together, these findings indicate that BSG depletion impairs ionocyte integration into the superficial epithelium without significantly changing the number of *Foxi1*-positive cells.

### BSG contributes to intracellular pH regulation during MCE development

Having established that loss of BSG.S results in intercalation defects in an ionocyte-specific manner, we investigated whether BSG.S knockdown also affected ionocyte function. Since Foxi1 is critical to proper differentiation of ionocytes from basal cells, embryos were injected with a Foxi1 morpholino (MO) to inhibit ionocyte differentiation. These embryos exhibited epidermal defects resembling those of BSG.S morphants (Fig. 3a). Ionocytes contribute to ion and pH homeostasis (*8*). To monitor intracellular acidity, we used pHrodo^TM^ Red–conjugated dextran, whose fluorescence increases as its local environment becomes more acidic (*18*). The probe was co-injected with membrane-GFP into a ventral animal blastomere at the eight-cell stage, and embryos were cultured to stage 28 for live imaging. pHrodo^TM^ fluorescence was detected throughout the developing epidermis with particularly strong signals in multiciliated cells (MCCs) and ionocytes, suggesting that the intracellular environment is more acidic in these cells than in other epidermal cell types (Fig. 3b). To investigate the contribution of ionocytes to intracellular pH regulation, we next assessed pHrodo^TM^ fluorescence following Foxi1 knockdown. Foxi1 morphants exhibited reduced pHrodo^TM^ fluorescence in MCCs (Fig. 3c). Similarly, BSG.S morphants showed substantially reduced pHrodo^TM^ fluorescence in both MCCs and ionocytes (Fig. 3d).

**Figure 3.**
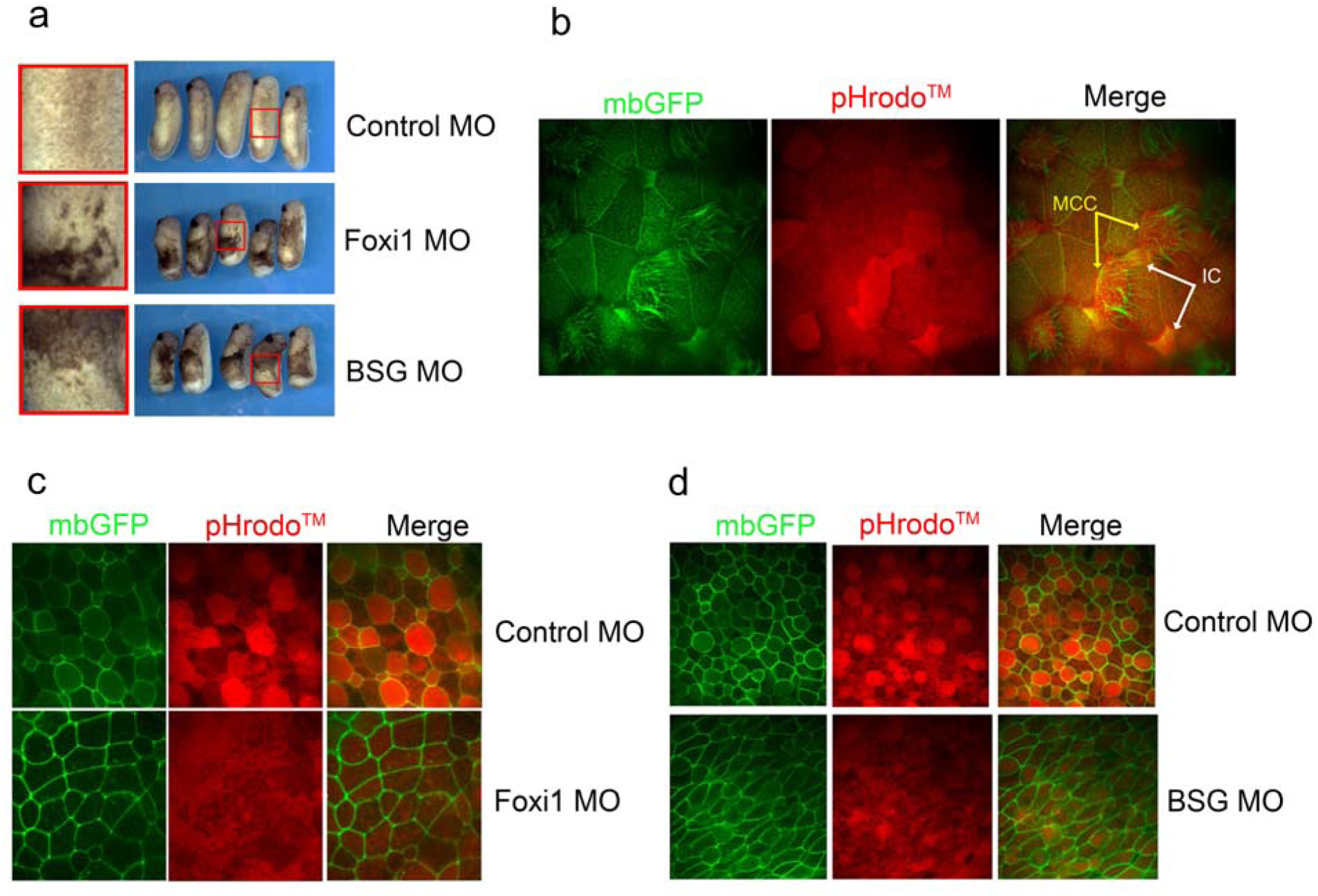
BSG regulates intracellular pH in the mucociliary epithelium. a. Representative images of stage 28 *Xenopus* embryos following injection of control MO, BSG.S MO, or Foxi1 MO (8 ng) into V.1 blastomeres at the 8-cell stage. Red boxes indicate regions shown at higher magnification. b. Confocal images of the MCE in wild-type embryos injected with pHrodo^TM^ Red Dextran^TM^ (red) and mbGFP (green). Arrows indicate MCCs (yellow) and ionocytes (white). c. Confocal images of pHrodo^TM^ Red Dextran fluorescence (red) in control and Foxi1 morphant embryos. d. Confocal images of pHrodo^TM^ Red Dextran fluorescence (red) in control and BSG.S morphant embryos.

Collectively, the similar changes in epidermal morphology and pHrodo^TM^ fluorescence following the loss of BSG.s and Foxi1 raise the possibility that BSG.S contributes to ionocyte-dependent pH regulation in MCCs during MCE development.

### BSG regulates epithelial cell-type composition during MCE development

Our previous results showed that BSG.S depletion reduced pHrodo^TM^ fluorescence and was associated with epidermal defects, with similar changes observed following Foxi1 knockdown. The MCE comprises MCCs, basal cells, goblet cells, small secretory cells, and ionocytes, whose coordinated activities support epithelial homeostasis (*4*). Ionocytes typically develop adjacent to MCCs, and Foxi1 knockdown depletes ionocytes and impairs cilia formation and basal body docking in neighboring MCCs (*5*). These findings support a non-cell-autonomous contribution of ionocytes to MCC development and function. We therefore examined whether BSG.S depletion also altered the other MCE populations. Compared with control MO-injected embryos, BSG.S knockdown increased the MCC population (Fig. 4a) and markedly reduced the population of basal cells as evidenced by the reduction of Tp63, a marker for basal cells (Fig. 4b). BSG.S overexpression did not significantly increase MCC abundance (Fig. 4c) but markedly increased the basal cell population (Fig. 4d). Similar with BSG.S knockdown, Foxi1 depletion increased the MCC population (Fig. 4e) and significantly reduced basal cell abundance (Fig. 4f).

**Figure 4.**
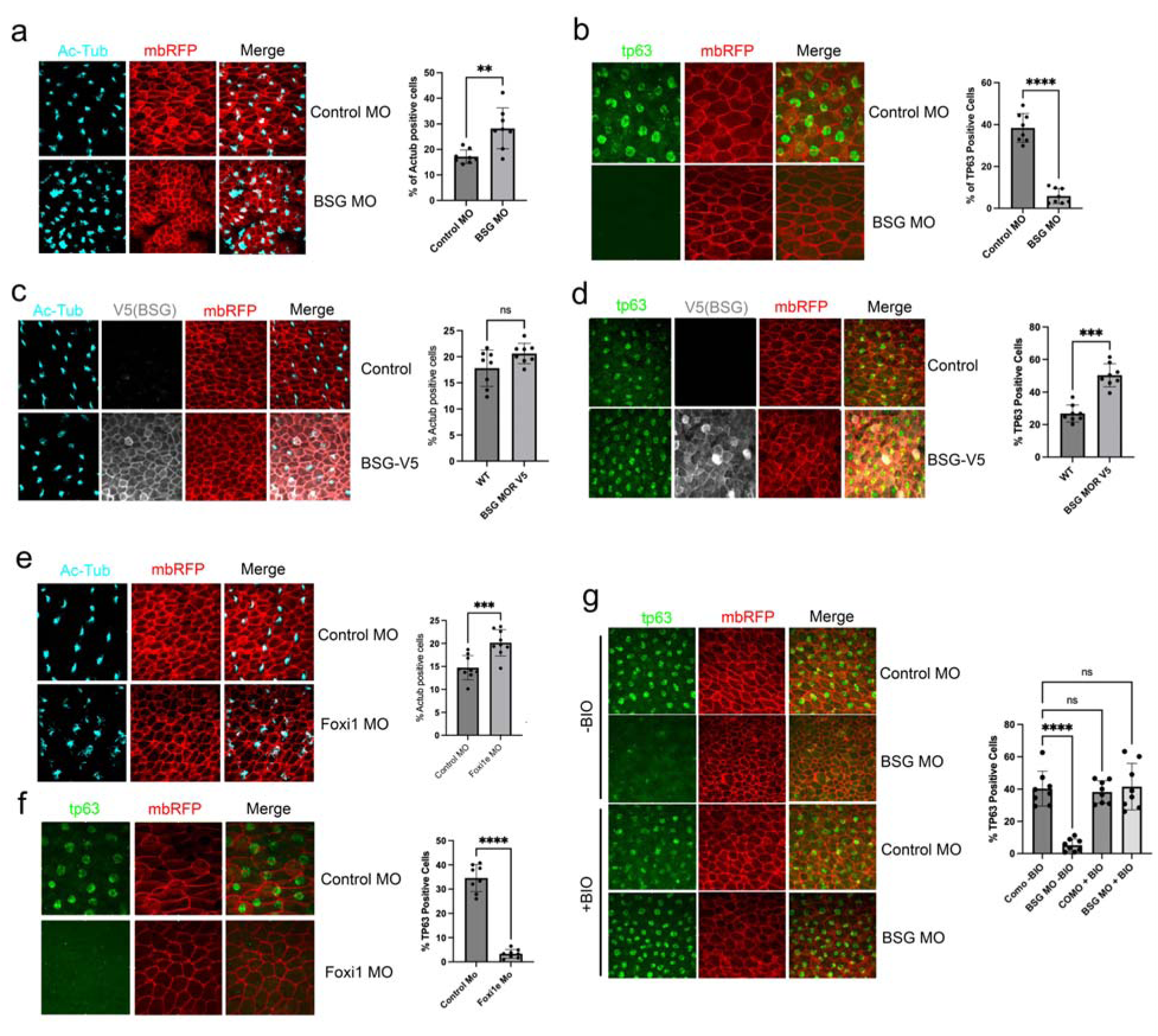
BSG.S modulates the balance of cell types in the developing mucociliary epithelium. a. Immunostaining for Multiciliated cells in control and BSG.S morphants (stage 28). control MO or BSG.S MO (8 ng/nl) into V.1 blastomere at the 8-cell stage. Acetylated tubulin (Ac-Tub, Cyan) is used to mark ciliated cells; mbRFP (red) marks cell boundaries. The graph shows the percentage of MCCs in the total cell population. b. Immunostaining for basal stem cells in control and BSG.S morphant embryos (stage 28). Tp63 (green) is used as a marker for basal stem cells. The graph shows the percentage of basal stem cells in the total cell population. c. Immunostaining for Multiciliated cells in control and BSG.S overexpressed embryos (stage 28). RNAs of BSG.S-V5 (300 pg) were injected into V.1 blastomere at the 8-cell stage. The graph shows the percentage of MCCs in the total cell population. d. Immunostaining for basal stem cells in control and BSG.S overexpressed embryos (stage 28). The graph shows the percentage of basal stem cells in the total cell population. e. Immunostaining for Multiciliated cells in control and Foxi1 morphants (stage 28). control MO or Foxi1 MO (8 ng/nl) into V.1 blastomere at the 8-cell stage. The graph shows the percentage of MCCs in the total cell population. f. Immunostaining for basal stem cells in control and Foxi1 morphant embryos (stage 28). The graph shows the percentage of basal stem cells in the total cell population. g. Immunostaining for basal stem cells in control and BSG.S morphants with or without BIO treatment from stage 13 to 28 (175μM). The graph shows the percentage of basal stem cells in the total cell population. Data are presented as mean ± [SD/SEM]. Statistical significance was determined using [paired t test and one-way ANOVA (Dunnett’s multiple comparisons test)]. *n* = [8] for each group. ns, not significant; \*\**p* < 0.001, \*\*\**p* < 0.0001, ****P<0.0001.

To explore mechanisms underlying the changes in MCE cell composition following BSG.S or Foxi1 knockdown, we investigated the potential involvement of Wnt/β-catenin signaling. This pathway promotes basal cell maintenance through ΔN-Tp63-dependent inhibition of epithelial differentiation (*19*). We therefore tested whether pharmacological activation of Wnt/β-catenin signaling could rescue the reduction in Tp63-positive basal cells associated with BSG.S depletion. Embryos were treated with BIO (6-bromoindirubin-3′-oxime), a GSK3 inhibitor, from stage 13 to stage 28 at a final concentration of 175μM. BSG.S knockdown markedly reduced the number of Tp63-positive basal cells, whereas BIO treatment largely restored this population (Fig. 4g).

Together, these findings support a role for BSG.S in establishing epithelial cell-type composition and suggest that Wnt/β-catenin signaling may contribute to the basal-cell phenotype associated with BSG.S depletion.

## Discussion

Ionocytes support mucociliary function by regulating the ionic environment in which mucus and motile cilia operate. In the *Xenopus* epidermis, ionocytes express proton pumps, carbonic anhydrases, and ion exchangers, including V-ATPase, CA12, and pendrin, that contribute to acid–base and electrolyte homeostasis (*8*). The influence of ionocytes also extends to neighboring MCCs. Ionocyte depletion through Foxi1 knockdown produces fewer and shorter cilia, abnormal basal-body positioning, and impaired ciliary beating. In mosaic experiments, MCCs adjacent to normal ionocytes exhibited rescue of ciliary morphology, supporting a local interaction between these cell populations, although the underlying mechanism remains unresolved (*5*).

Our loss-of-function analysis identifies BSG.S as a regulator of ionocyte radial intercalation. BSG.S depletion impaired ionocyte integration into the superficial epithelium without altering the number of Foxi1-positive cells. These findings suggest that BSG.S contributes to the morphogenesis and integration of specified ionocytes. One possible mechanism involves extracellular matrix (ECM) remodeling, as BSG has been linked to matrix metalloproteinase (MMP) production and matrix invasion in tumor–stromal systems (*10*). However, our analysis of the single-cell RNA-sequencing dataset did not identify specific MMP induction during MCE development. This observation does not exclude changes in protease localization or activity. Direct analysis of ECM organization and MMP activity will be needed to determine whether matrix remodeling contributes to the intercalation phenotype.

Our live-imaging experiments also revealed differences in pHrodo^TM^ fluorescence among epithelial cell types. Under control conditions, MCCs and ionocytes exhibited stronger fluorescence than goblet cells, consistent with more acidic intracellular environments (*20, 21*). BSG.S depletion reduced this fluorescence in MCCs and ionocytes, suggesting reduced intracellular acidity. Foxi1 depletion similarly reduced the signal in MCCs, raising the possibility that ionocytes influence pH regulation in neighboring cells. The physiological significance of this fluorescence pattern remains unclear, including whether it contributes to MCC maturation or reflects the functions of differentiated cells. Calibrated measurements with defined probe localization will be important for distinguishing changes in cytosolic pH from changes within intracellular compartments.

BSG.S depletion also altered epithelial cell-type composition, increasing MCC population and reducing basal cell population. Similar changes following Foxi1 depletion suggest a potential relationship between ionocyte development and the establishment of other epithelial cell populations. Wnt/β-catenin signaling regulates basal cell maintenance through ΔN-Tp63-dependent inhibition of differentiation (*19*). In our experiments, treatment with BIO, a GSK3 inhibitor used to activate Wnt/β-catenin signaling, restored the number of Tp63-positive basal cells in BSG-depleted embryos. This rescue supports possible involvement of Wnt/β-catenin signaling in the basal cell phenotype, although it does not establish that BSG.S directly regulates the pathway. A potential connection between BSG-associated pH changes and Wnt signaling involves the pH-dependent stability of β-catenin. Previous studies in mammalian cells and Drosophila demonstrated that elevated intracellular pH increases β-catenin binding to the E3 ubiquitin ligase β-TrCP, promoting its degradation. This pH sensitivity depends on the conserved His36 residue within the β-TrCP recognition motif and operates together with phosphorylation of the β-catenin degradation motif. Deprotonation of His36 at higher pH favors β-TrCP association, providing a mechanism through which intracellular alkalinization can reduce β-catenin stability (*22*). These observations suggest a testable model in which BSG.S loss elevates intracellular pH, promotes β-catenin degradation and weakens signaling that supports basal cell maintenance. Such a mechanism could contribute to the loss of basal stem cells and increase in MCCs following BSG.S depletion. Future studies and development of new tools will be required to determine cell-type specific levels of cytosolic pH, β- catenin stability and Wnt transcriptional activity.

Our study identifies an important role for BSG.S in regulating ionocyte position and function, demonstrating the importance of ionocytes in maintaining homeostasis and cellular composition within the mucociliary epithelium during embryonic development.

## Material and methods

### Xenopus laevis

Wild-type *Xenopus laevis* were obtained from Nasco, USA. All animal procedures reported in this study that were performed by NCI-CCR affiliated staff were approved by the NCI Animal Care and Use Committee (ACUC) and in accordance with federal regulatory requirements and standards. All components of the intramural NIH ACU program are accredited by AAALAC International.

### Oligos, plasmids, reagent, and mRNA synthesis

The cDNA clone that encoded full-length *Xenopus* BSG.S was obtained from Source BioScience (Gene ID: 734661). Various deletion mutants were generated in the pCS107 vector using the QuikChange II Site-Directed Mutagenesis Kit. Capped sense RNAs were transcribed using the mMessage mMachine SP6 kit (ThermoFisher) after linearization with ASP718. The sequences of MOs are as follows; BSG.S MO: 5’-AAGCAGCCCGAGACCCATTTTCCTG-3’; Foxi1 MO: 5’- ATAATTGCCCTTGCCTGTAAAGAAAA -3’. To assess intracellular acidity, 100 pg of pHrodo™ Red– conjugated dextran (P10361, Invitrogen) was injected into the V1 blastomere of each embryo at the 8-cell stage. The Pendrin–GFP DNA construct was a gift from Dr. Brian Mitchell at Northwestern University.

### Hybridization chain reaction RNA fluorescence in situ hybridization (HCR RNA-FISH)

HCR probe sets were designed by Molecular Instruments. *Xenopus* embryos were collected at stage 24 and then processed for HCR following the HCR v3.0 protocol for whole-mount zebrafish embryos. For the cross-section, Embryos were embedded in 4% low-melting agarose gel (2070-OP, Sigma) and were sectioned with a thickness of 100 μm with the vibratome (LEICA VT 1200 S). HCR images were taken by confocal microscopy (Zeiss LSM880).

### Immunofluorescence and confocal microscopy

immunofluorescence microscopy was carried out using standard protocols. Briefly, the embryos were fixed within MEMFA (4% formaldehyde in 1× MEM salt) at 4 °C for overnight and then dehydrated with 100% methanol. The following primary antibodies were incubated after blocking with the filtered 10% goat serum in 1x PBS: Rabbit anti-tp63 (1:1,000, ab735, Abcam), Mouse anti- SLC4A1 (AE1) (1:100, IVF12, DSHB), Mouse anti-acetylated tubulin (1:1000, T7451, Sigma-Aldrich), Mouse anti-V5-Alexa Fluor-488/555/647 (1:500, 2F11F7, ThermoFisher), and Rabbit anti-GFP-Alexa Fluor-488 (1:500, A- 21311, ThermoFisher). The secondary antibodies used were Alexa Fluor-488 or Alexa Fluor-594 conjugated Goat anti-rabbit IgG or anti-mouse IgG (1:500, Invitrogen). For F-actin staining, Alexa Fluor™ 488 Phalloidin (A12379, ThermoFisher). The samples were mounted and imaged using Zeiss LSM880 with Airyscan laser scanning confocal microscope.

### Whole-mount in situ hybridization (WISH)

*Xenopus* embryos were collected at stage 16 or 24 for hybridization with the Foxi1 or BSG.S probes. Embryos were injected with membrane-RFP and various mRNAs or MOs to distinguish the injected side of embryos. The embryos were then processed for whole-mount *in situ* hybridization using standard methods.

### Reverse transcription and quantitative real-time PCR

Reverse transcription was carried out by using a SuperScript™ IV First-Strand Synthesis System (ThermoFisher). The PCR reactions were performed with SYBR™ Green PCR Master Mix (4309155, Applied Biosystems™) using QuantStudio™ 5 Real-Time PCR System (A34322, Applied Biosystems™). ODC was used for normalization.

### Statistics and reproducibility

The pilot experiments were conducted for all experiments to evaluate RNA and MO dosage, and then full-scale experiments were performed at least three independent times. All experiments were performed blinded with an order of testing randomized. The sample size was determined as indicated in the figures and the specific statistical method was not used. Dead cells and embryos were excluded from all experimental analysis. Embryos that have mistargeted injection also were excluded before analysis (the target injection was confirmed by co-injection of lineage tracer (membrane-GFP/RFP RNAs). ImageJ program was used for all quantification. Normality of data was tested using Kolmogorov–Smirnov’s test, D’Agostino and Pearson omnibus normality test and Shapiro–Wilk normality test using Prism 10. The data were considered as normal if found as normal by all three tests. Data sets following a normal distribution was compared with Student’s t-test (two-tailed, unequal variances) or a one-way analysis of variance (ANOVA) with Dunnett’s multiple comparisons post-test in Prism10. The data that did not follow a normal distribution were compared using Mann–Whitney’s test or a non-parametric ANOVA (Kruskal–Wallis with Dunn’s multiple comparisons post-test) using Prism 10. Cross comparisons were performed only if the overall p-value of the ANOVA was <0.05.; error bars: SEM.

## Acknowledgements

We thank Dr. Chris Westlake for technical support for microscopy, and all members of the Cancer and Developmental Biology Laboratory for discussion and comments. We are grateful to the Francisco Perieria Lobo for single-cell-RNA-seq analysis. This research was supported by the National Cancer Institute, National Institutes of Health, Intramural Research Program, and was funded in whole or in part with federal funds from the National Cancer Institute, National Institutes of Health under contract HHSN26120080001E. The content of this publication does not necessarily reflect the views or policies of the Department of Health and Human Services, nor does mention of trade names, commercial products, or organizations imply endorsement by the U.S. Government.

